# Methylation at CpG sites in genomes of aphid *Acyrtosiphon pisum* and its endosymbiont *Buchnera*

**DOI:** 10.1101/2021.08.26.457790

**Authors:** Mathilde Clément, Martine Da Rocha, Sandra Agnel, Guenter Raddatz, Alain Robichon, Marc Bailly-Bechet

## Abstract

Pea aphid *Acyrtosiphon pisum*, a sap-feeding insect, has established a mutualistic relationship with an endosymbiotic bacteria (*Buchnera aphidicola*) that constitutes an evolutionary successful symbiosis to synthetize complex chemical compounds from a nutrient deprived diet. In this study, led by the presence of DNMT1 and a putative DNMT3 methylase in the aphid genome, we investigated the distribution of the methyl groups on 5’cytosine in CpG motifs on the whole genomes of host and endosymbiont, and looked into their correlation with gene expression. The DNA methylation turned to be present at low level in aphid (around 3% of total genomic cytosine) compared to mammals and plants, but increased to ∼9% in genes. Interestingly, the reduced genome of the endosymbiont *Buchnera* also shows global low level of methyl cytosine despite the fact that its genome does not shelter any DNA methylase. This finding argues for the import of DNA methylase from the host to the endosymbiont. The observed differences in methylation patterns between two clonal variants (host plus endosymbiont) are reported along with the differences in their transcriptome profiles. Our data allowed to decipher a dynamic combinatorial DNA methylation and epigenetic cross talk between host and symbiont in a clonality context that might count for the aphid adaptation to environment.

## Introduction

Cytosine methylation represents an epigenetic modification of the genome widespread in eukaryotic world from plants, mammals to insects and in prokaryotic organisms [1,2,3,4]. In contrast, DNA methylation has been lost in the budding yeast *Saccharomyces cerevisiae* and the nematode worm *Caenorhabditis elegans* [5]. Generally, the cytosine methylation appears highly enriched in repetitive DNA like centromeric regions and transposons [1,6]. The family of conserved cytosine methyltransferases (DNMTs) is responsible for the addition of methyl group at CpG sites and their deletion in mammals leads to lethality, which indicates a major still elusive role in development [7,8,9]. The patterns of added methyl group on bases, mainly cytosine, have been found highly dynamic in most of species from different orders [8,10]. To this respect, DNA methylation undergoes large transitions during early embryonic development in mammals [7,9], regulates the queen/workers phenotypes in honeybee [11,12] and targets DNA cleavage of invading phages in bacteria by directing restriction enzymes [13]. On the other side, several factors influence DNA methylation like the TET proteins responsible for demethylation through successive oxidative steps, leading to 5-hydroxymethylcytosine (5hmC), 5-formylcytosine (5fC) and 5-carboxylcytosine (5caC) [14]. Both 5fC and 5caC are excised and repaired to regenerate unmodified cytosines by the concerted action of thymine DNA glycosylase (TDG) and the base excision repair (BER) enzymes [15,16]. The current accepted dogma is that the CpG dinucleotide remains the primary site for DNA methylation, but there is an emerging evidence for non-CpG methylation in cytosine and/or adenine in *Drosophila* [17], *C. elegans* [18], *Chlamydomonas* [19] and human cells [20,21,22]. In insects, many gene body sequences and promoter regions show a variable level of methylation at remarkably lower level ranges compared to those reported in plants and mammals [5,6,8]. The function of these DNA methylations, regarding their implication in gene regulation, is debated with a general view that they might have apparently opposite roles. For instance, DNA methylation in non-promoter intergenic regions and gene bodies is known to antagonize the Polycomb mediated gene repression, opening the way for the maintenance of active chromatin state and gene expression in the vicinity [23]. On the other side, DNA methylation has a known repressive role in silencing transposons elements and also in preventing transcriptional machinery to operate on many genes [5,6,8].

In this report, we explored the DNA methylation of a clonal species of insect, the aphid *Acyrtosiphon pisum*, in order to assess its role in adaptation to fluctuating environment. Aphids are endowed of polyphenism, which means that morphological changes like winged or wingless are produced depending on environmental conditions like density of population [24,25,26,27]. The reproductive modes can change also giving viviparous parthenogenesis and sexual forms in response to the photoperiodicity and temperature factors [28,29]. Under reduced light conditions in autumn, sexual morphs (males and oviparous females) are produced by parthenogenetic mother and this shift of reproduction mode seems to be regulated by juvenile hormone (JH) titers in the mothers [28,29]. Therefore, aphids exhibit both sexual and asexual reproduction depending on the season: diapausing eggs for overwintering and nymphs produced by clonality in spring/summer, which is an evolutionary success conferring their survival over 250 millions years [28,29]. As JH has been reported to be methylated [30], this regulation may may be related to DNA methylation. Another adaptive phenotype in aphids is the methylation of a gene coding for an esterase in an aphid species (*Myzus persicae*), which has been described as essential for its upregulation and leads to pesticides resistance [31,32,33].

Furthermore, the aphid *A. pisum* uses the obligate endosymbiont *Buchnera* to synthetize essential compounds for which it lacks the enzymatic equipment to produce them [34,35,36]. The endosymbiont *Buchnera* is a proteobacteria that has a highly reduced genome compared to the most related bacteria, *E. coli*, and is characterized by the loss of key enzymes involved in carbohydrate and lipid metabolizing pathways [37]. Furthermore genes coding DNA methylases are absent in its genome [37]. In contrast, *Buchnera* produces many essential amino acids like tryptophan, vitamins and lipids that are not present in the plant phloem and that the host is not able to synthetize [37]. A phylogenetic analysis led to the conclusion that the symbiotic relationship started 200–250 millions ago and the DNA sequencing reveals that *Buchnera* chromosome is 640,681 base pairs (bp), one of the smallest of the completely sequenced bacterial genomes, plus two circular plasmids (around 8,000 bp) [37].

In this report, two clonal variants of aphids raised at different temperatures (9°C and 20°C) and originated from the same founder mother were investigated at wide scale genomic level in order to unravel putatively non-mendelian mechanisms to transmit phenotypic traits. We focused on the methyl cytosine analysis due to technical advances that allow to determine their precise genomic location, in contrast with modified adenine. In this study, the whole genome bisulfite sequencing (WGBS) was performed to establish the complete methyl CpG mapping. Interestingly, we show that without harboring methylase genes, the endosymbiont genome is methylated. The differential transcriptome analysis was also performed in parallel with the methylome mapping between the two variants (selected by temperature) for both the endosymbiont and aphid host. A comprehensive highly methylated gene list in aphid is presented; those highly methylated genes mostly show a high expression. Moreover our analysis highlights the existence of hot spots of methylation in both the host and endosymbiont genomes. As an example, an aphid contig presenting a clusterization of genes related to the transcriptional factor gene *lola* in *Drosophila* turned out to be highly methylated. Finally, our data argue in favor of host methylase diffusion/transport inside the endosymbionts to operate in situ DNA methylation although its role remains elusive.

## Results

### Pipeline of analyses

The methyl CpG mapping and quantification was conducted in the two clonal variant genomes (aphid and endosymbiont) selected by temperature and originated from the same parthenogenetic mother ancestor. The aim of this study was to determine a precise map of methyl CpG in both host and endosymbiont genomes and whether this map diverges/overlaps between the *orange* (temperate adapted) and *green* (cold adapted) variants. In parallel, RNA-seq analyses were carried out in order to assess whether 1) a panel of genes is specifically up or down regulated in *orange* or *green* host/endosymbiont variants, 2) a correlation exists between the degree of DNA methylation of genes (gene bodies and promoters) and their expression level. The scheme of the overall procedure is described in Fig. 1. Raw numbers obtained by Bismarck analysis are shown in Supp. Table S1.

**Figure 1:**
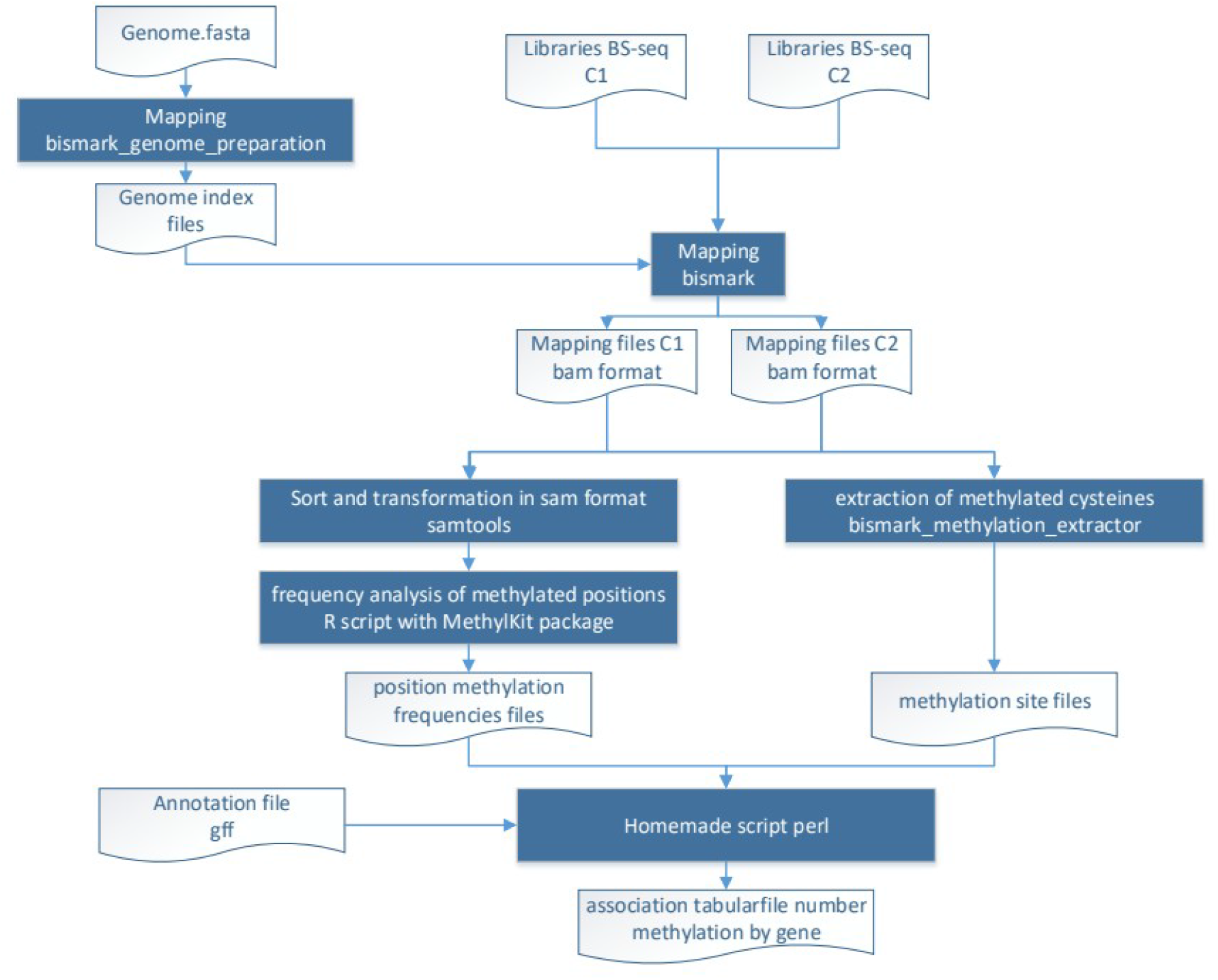
Pipeline of analysis.

### The aphid methylome and transcriptome

First, we characterized the patterns of methylation across the genome of *A. pisum*. Methylation in insects is predominantly at CpG motifs rather than CHG or CHH sites (h= A,C, or T) commonly observed in bacteria, and we confirm it in our data (See Supp Table S1). The conserved DNA methylase genes DNMT1 and DNMT3 present in the *Acyrthosiphon pisum* aphid are responsible for these additions. Globally, 3.4% of the A. pisum genome is methylated (see Supp. Table S1). Those methylations are targeted on genes: of the estimated 6.83 10^6^ CpG sites in *A. pisum* genes (including introns, 5’ and 3’ UTR), 0.94 10^6^ were identified by bisulfite sequencing; out of those, 82863 were methylated, giving an average methylation estimate of 8.79% (8.65% in the *green* strain and 9.05% in the *orange* one). A strong correlation between methylation in *green* and *orange* variants (r=0.8) demonstrates that methylation patterns are not random (Fig. 2A). Moreover, we observed that a high methylation level corresponds to a medium to high FPKM in the RNAseq determination (Fig. 2B), leading to a positive correlation (Spearman rho=0.33, p<10^−16^) between methylation and expression. The WGBS analysis of the aphid genome reveals unambiguously that two major groups of genes are easily observable: a subset of the genes that turned out to be highly methylated, whereas the other major part of total genes was not methylated at all (Fig. 2B). These two populations of genes are similar in the two clonal variants (Fig. 2A), which indicates that this binary distribution is preserved within the clonal variants selected by environmental conditions.

**Fig 2:**
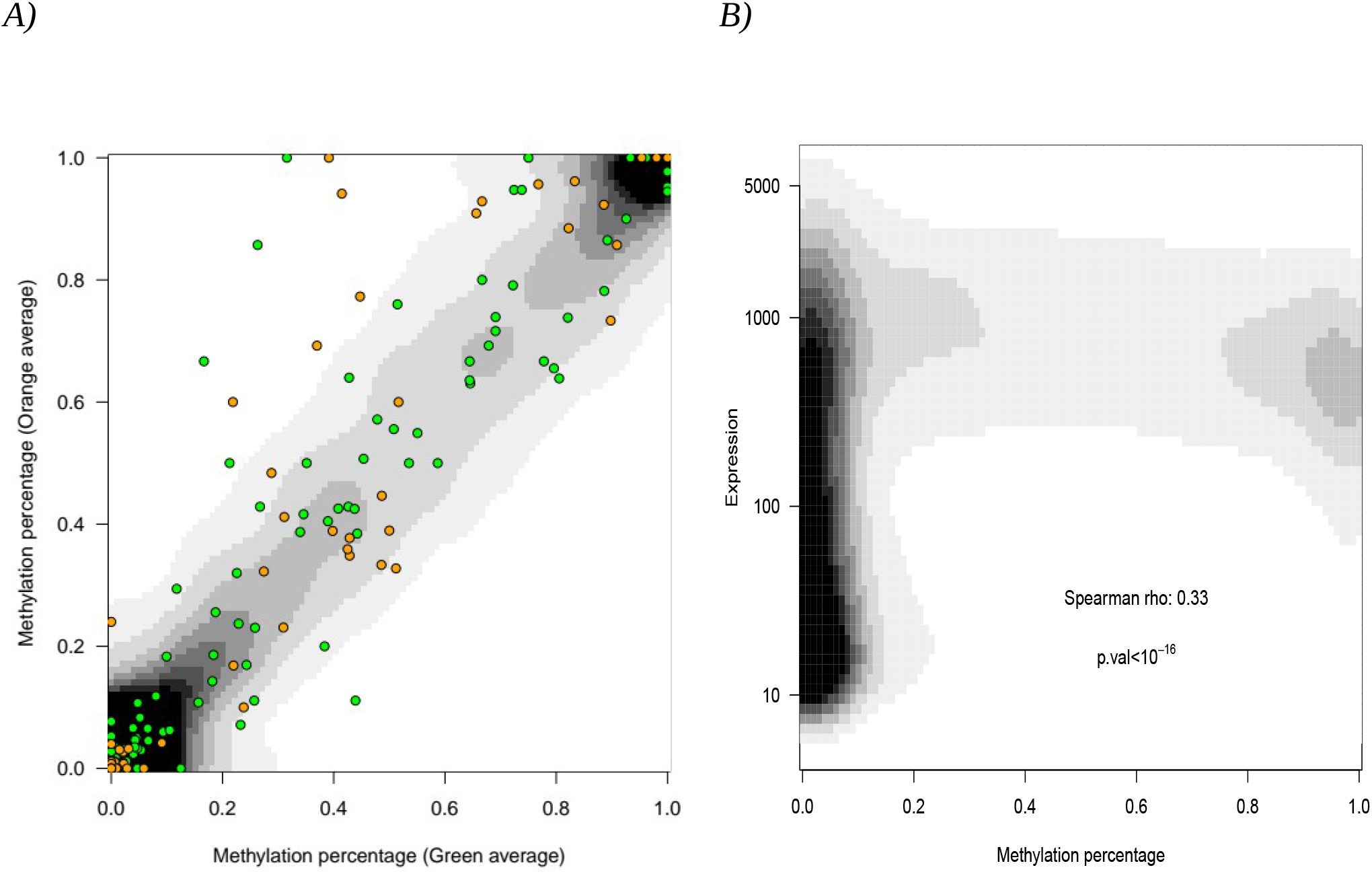
Correlation between methylated genes in green and orange aphid clonal variants and global correlation expression/methylation A) x-axis: methylation percentage in the “green” aphid strain. y-axis: methylation percentage in the “orange” aphid strain. Instead of representing each gene as a point, results are shown using a density map, darker shade indicating the presence of more genes. Note the high gene density on the diagonal. Green dots are overexpressed in the “green” strain relative to the “orange” one, orange dots are overexpressed in the “orange” strain relative to the “green” one. B) Methylation percentage (x-axis) vs Expression (y-axis, log scale) in aphid (both “orange” and “green” variants). Instead of representing each gene as a point, results are shown using a density map, darker shade indicating the presence of more genes. Correlation coefficient and the corresponding p-value are shown. NB: methylation is measured on the full gene, including non coding parts such as intron and 5’/3’UTR.

We could not compare precisely our results to those obtained in [38], the methylated reads obtained in this work being too long relative to the accuracy of the present WGBS analysis, and global correlations showing non significant overlaps. However, we could compare our results to those obtained previously in [30], who identified 12 methylated genes. Out of those 12, we could identify 10, and 5 out of those 10 were found methylated in our study (Supp. Table S2), with a methylation percentage above average (Wilcoxon test, p<0.007); no signal, neither high or low, was obtained for the 5 other ones. We hypothesize this is due to the random sampling of sequences consequent to the bisulfite sequencing methodology.

Comparison of the level CpG methylation in gene subparts plus promoters revealed that exons and 3’UTR CpGs are more methylated (16.6 and 18.6% respectively) than introns (5.8%), 5’UTR (5 %) or promoters (5.5 %) (Fig. 3). This trend is also visible when looking at the distribution of methylation, which is U-shaped for all categories with a higher percentage of sequences methylated at more than 95% in exons and 3’UTR than other categories (Fig. 3). Finally, considering the global population of genes, the methylation between different subparts of genes is positively correlated (Table 1) which means that highly methylated genes will be significantly methylated in all their subparts. This argues for the paradigm of an “all or none” methylation pattern acting globally on genes, and then observed concomitantly in all the subparts of genes. This might suggest that the methylation process is regulated at a regional level in genome instead of recognizing specific sequence cues for its targeting. At a finer scale, we can see that correlations are higher, and then methylation more homogenous, between the introns/exons/3’UTR subparts, i.e all subparts located after the start codon.

**Figure 3:**
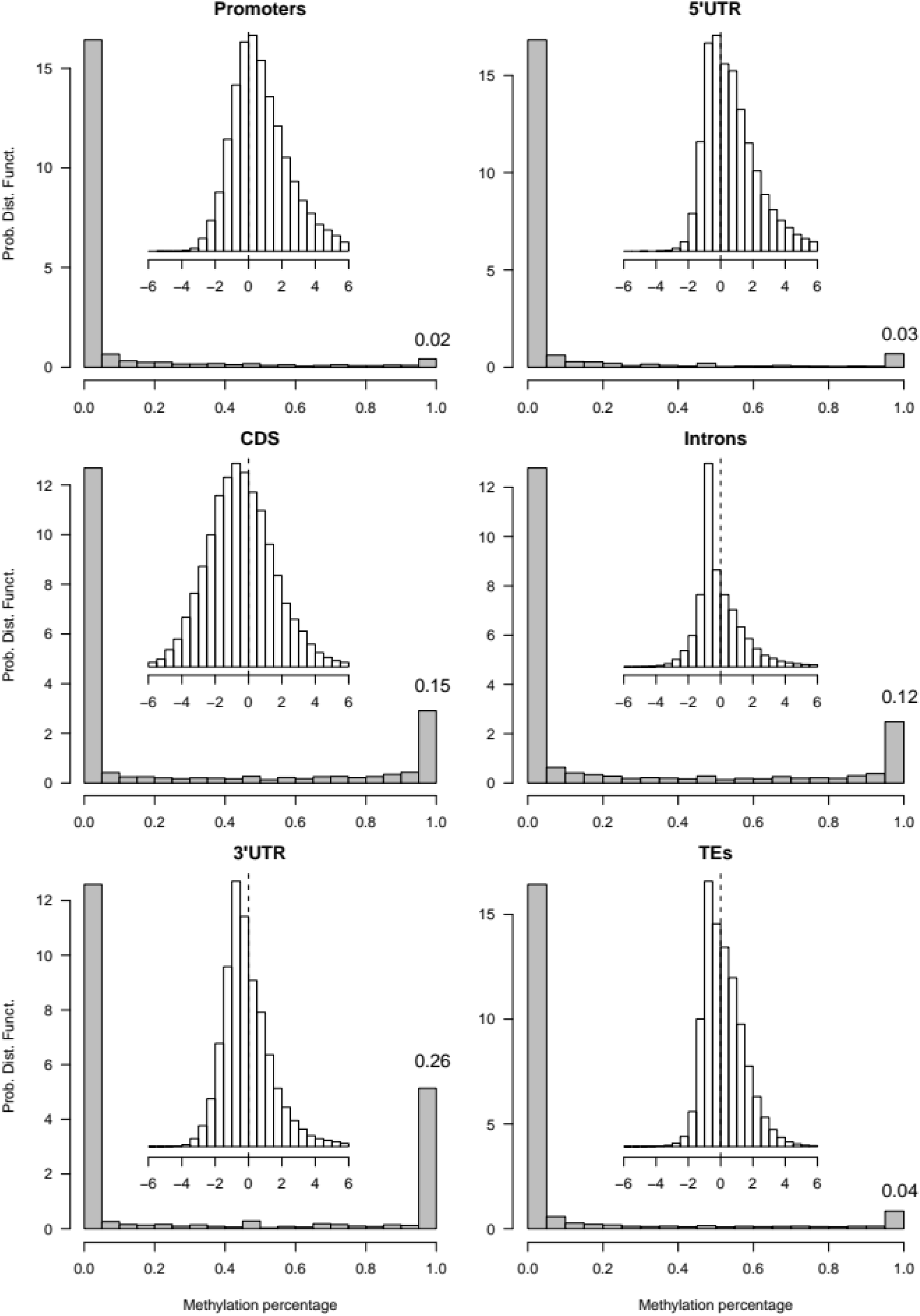
Methylation percentage distribution among subparts of gene sequences (namely: exons, introns, promoters, 3’UTR, 5’UTR) and for TE sequences. In grey, the large histogram shows the distribution of methylation percentage among sequences. x-axis indicates methylation percentage aned y-axis probability density function. The bar corresponding to 1 on the x-axis indicates a methylation percentage above 0.95, with percentage of the sequences concerned written above. In inset, the white histogram shows the distribution of the CpG dinucleotide enrichment (measured as a z-score, see Methods) for each category. Dotted line indicates a 0 level enrichment, i.e. random CpG presence/absence relative to the subpart G+C content.

**Table 1:**
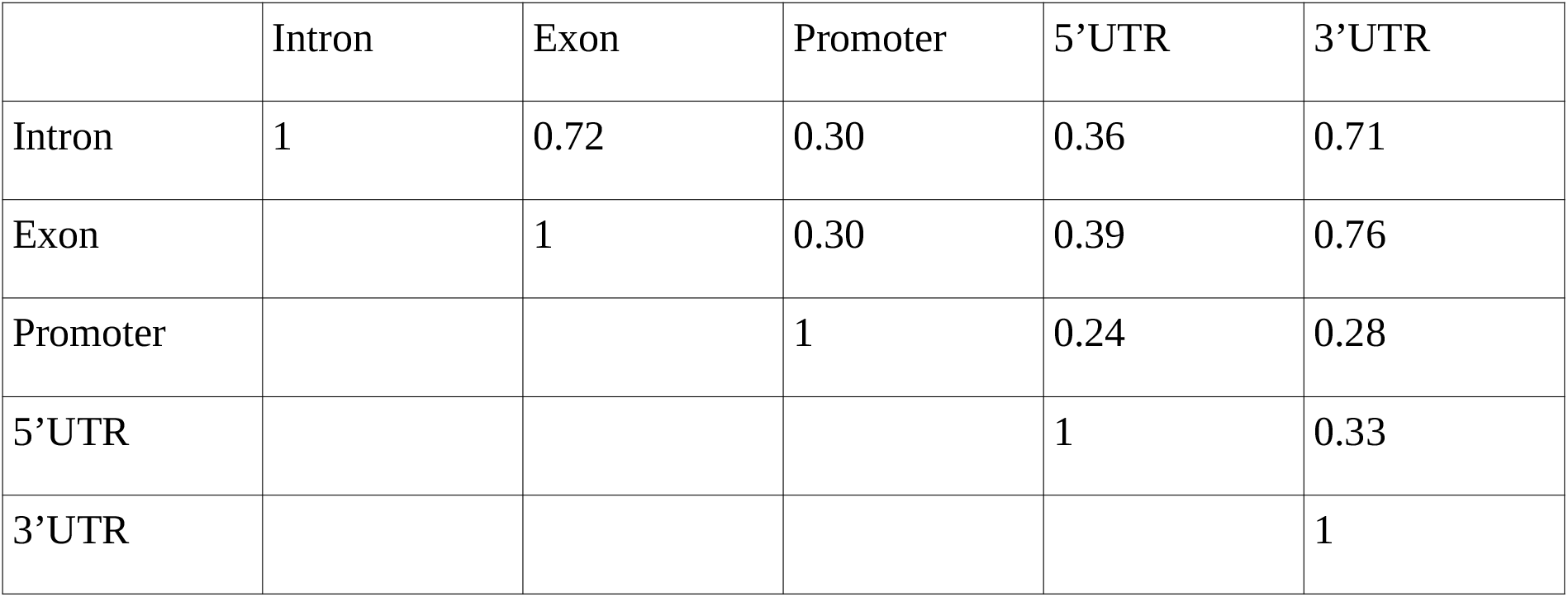
Intragenic correlation (Spearman Rho) between methylation percentages of genetic sequences subparts.

On the other side, transposable elements (TE) show a methylation average of 7.1%, which is closer to non-coding sequences than to exons. For TEs, we also observed a U-shaped distribution regarding the percentage of methylation, which seems to indicate that TEs are not evenly methylated but, as other genes and genes subparts, show a strong heterogeneity: few are heavily methylated and the majority is not at all (Fig. 3).

We then investigated whether those methylation patterns (as percentages of methylated CpG) could be related to the distribution of CpG dinucleotides among genes subparts. We computed the enrichment z-score [39] of CpG dinucleotides for all sequences relative to the global genomic content of the subpart, and show its distribution (Fig. 3, inset histograms). This score was preferred to the rho factor because it is not affected by sequence length. In all the subparts except promoters and 5’UTR, a peak for negative enrichment, i.e. counter selection of CpG dinucleotides relative to the subpart G+C content, was observed. Promoters and 5’UTR are also the subparts where less methylation is seen. Methylation spots hence appear related to a counter selection on CpG dinucleotides, seemingly starting at the start codon and coupled with a higher – and more homogeneous -- methylation on exons, introns and 3’UTR.

The transcriptome analysis of the two aphid host variants showed a panel of significantly modified genes. The analysis revealed that 91 genes were significantly differentially expressed, the complete list of which is provided in Table S3. We noticed that many of them are uncharacterized proteins or show hypothetical functions based on sequence conservation. We observed that the differentially expressed genes with strong expression in the *orange* variant showed no significant GO enrichment, maybe due to the small number (19) of genes concerned. On the contrary, the *green* variant appeared enriched in genes related to neuron development and differentiation, and microtubule cytoskeleton organization.

Although unambiguous hotspots of methylation were observed in the genomes of both variants, we could find no evidence of a link between differential expression and differential methylation (Spearman correlation test, rho=0.11, p=0.3) on genes differentially expressed.

Regarding the hotspot of methylation in aphid genome, we identify 912 genes 100% percent methylated and being long enough so that this total methylation can be considered as non random given the average methylation level in the genome (see Table S4 for the identification of each gene). The GO term enrichment of these genes emphasizes cellular protein metabolic process (corrected p<0.0261) and cellular localization (corrected p<0.0862) in organelles (corrected p<0.024); terms related to Golgi organization and transport from Golgi to the membrane are non significant but show p-values smaller than 0.15, hinting that maybe a subpart of those genes may be related to these processes (Full data in Supp. Table S5).

Interestingly, those 912 hypermethylated genes show a clear tendency to be counter-selected for CpG dinucleotides, relative to the whole genome (t-test on z-scores of CpG enrichment, hypermethylated genes average -2.32, other genes average 0.42, p<2.10^−16^), confirming our previous hypothesis about counter-selection of CpG in methylated regions. Moreover, those 912 genes are not distributed evenly among the aphid genome scaffolds, but tend to be concentrated on some scaffolds (permutation test, p<10^−3^). When looking at the distribution of those hypermethylated genes in the aphid genome, we find a large concentration of them on a single contig where 33 out of 57 genes turned out to be hypermethylated. Amazingly half of these genes (16 out of 33) shares structural features: they are related to longitudinal lacking protein (Lola) first described in *Drosophila* that acts on ubiquitous developmental stages, oogenesis and spermatogenesis, neural wiring and eye development [40,41,42]. This molecule belongs to zinc finger C2H2 superfamily DNA-binding transcription factor [43]. This C2H2 motif can be arranged in clusters presenting multiple adjacent C2H2 or in separated paired C2H2 and also in triple C2H2 depending on proteins that harbor them [44]. If the main function of these proteins is to bind DNA through their zinc finger domains, some of them can also bind RNA, protein and lipids in relation with their C2H2 configuration [45, 46]. These structural features explain why the superfamily of proteins harboring C2H2 motifs exhibit ubiquitous functions: for instance *lola* is known to be involved in *Notch* signaling, in targeting axons and dentrites and in controling the retrotransposon dispersion [40,41,42]. Finally, 4 of the genes not related to Lola are zinc-finger proteins. Therefore the 20 hypermethylated genes, all of them zinc fingers and most related to *lola* genes, argue in favor of a transcriptional activity in early developmental stages under control of DNA methylases. In aphid (*A*.*pisum*) *lola* family of genes are co-localized on the same short contig indicating that the regional clusterization is determinant and pivotal for methylase action.

### The endosymbiont methylome and transcriptome

We decided to look at the CpG methylation in the endosymbiont *B*.*aphidicola*. Interestingly, even if the endosymbiont does not harbor genes encoding for DNA methylases, the numbers of mCpGs detected was 1746 (9.88%) and 2366 (13.4%) in the *orange* and the *green* variant, respectively on a total of ∼17650 CpG. Those methylation levels are in the same range as those of the host. These values are not random, as we observe a low yet significant correlation at the gene scale between the 2 replicates for each clonal variant (Spearman rho=0.26, p<2 10^−10^ for the orange variant and rho=0.12, p<0.0032 for the green variant). The *Buchnera* genome methylation was verified by manual bisulfite sequencing performed on 6 genes found methylated in WGBS (see Table S6). The two experimentations gave similar results arguing for the robustness of the finding and supporting the reality of *Buchnera* genome methylation at CpG sites. This supports the paradigm of cross methylation between host and endosymbiont genomes.

Comparing both clonal variants, we observe significant correlations between them (Fig. 4A), which indicates that methylation may not be highly dynamic in them. The main feature distinguishing the methylation patterns in *Buchnera* from those of its aphid host is the absence of sequences fully methylated.

**Fig 4 :**
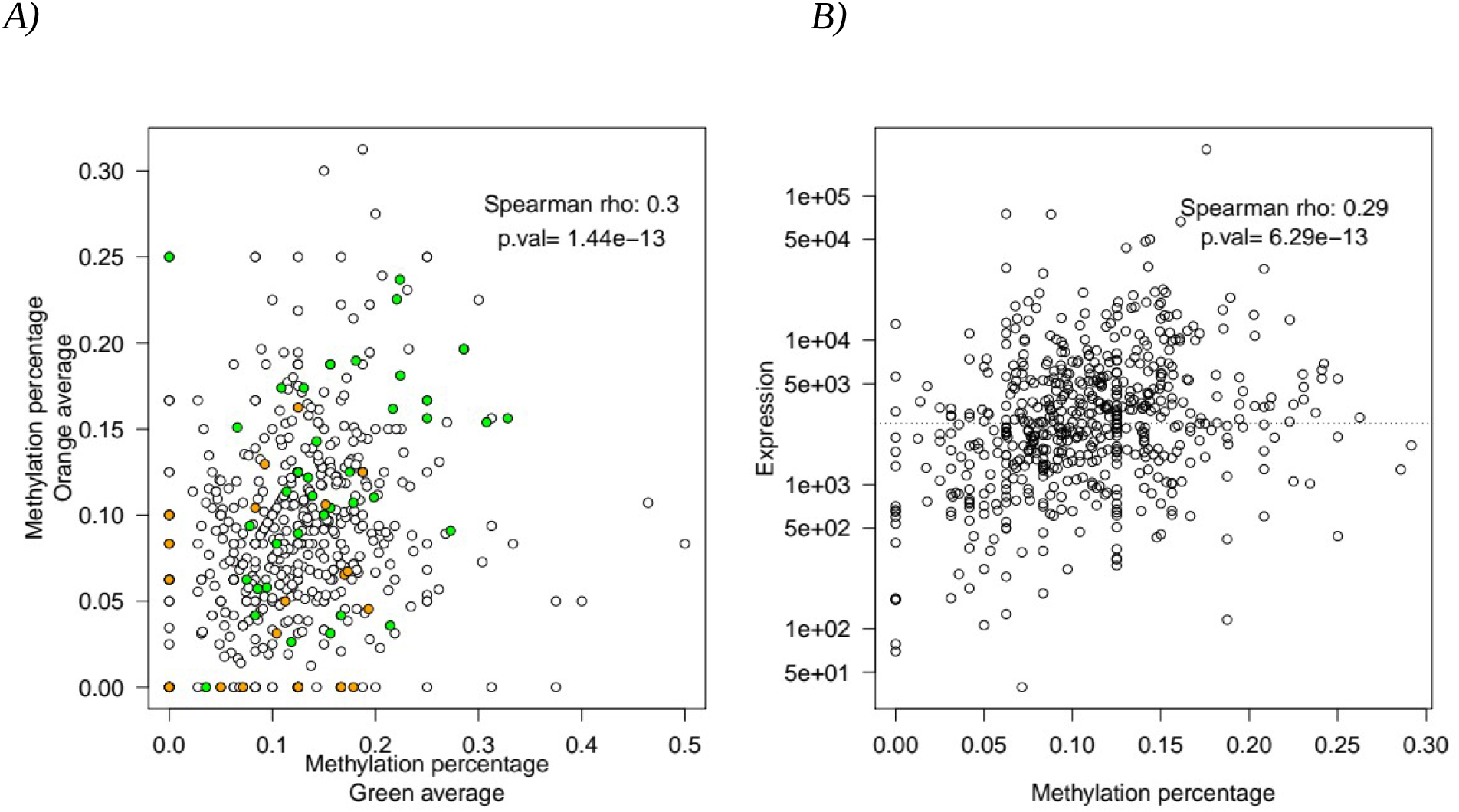
Intensity in Buchnera gene methylation: Correlation between Buchnera originated from green and orange variants and correlation between levels of methylation and expression A) x-axis: methylation percentage in the “green” Buchnera strain. y-axis: methylation percentage in the “orange” Buchnera strain. Each dot is a gene; green dots are overexpressed in the “green” strain relative to the “orange” one, orange dots are overexpressed in the “orange” strain relative to the “green” one. B) Methylation percentage (x-axis) vs Expression, measured as DeSeq2 BaseMean (y-axis, log scale) in Buchnera. Black circles are individual genes Spearman correlation coefficient and the corresponding p-value are shown. The dotted line corresponds to the average of levels of expression for the full genome.

The analysis showed also that for a large panel of genes, the difference of methylation between the two endosymbionts was only quantitative: methyl marks were found in genes in both *green* or *orange* variants for a vast majority of them. In contrast methyl marks found exclusively in either *green* or *orange* variants concerned a restrained panel of genes (Fig 4A). This indicates that a “all or none” rule was not observed in those genomes, for neither variant.

We also looked at the link between methylation and expression. As for the aphid host, we find a positive correlation between expression and methylation (Fig. 4B), which is still true for both clones studied separately. We also did not find any link between differences in methylation between clones and differences in expression between clones (Spearman correlation test, rho=0.046, p=0.26). The RNA seq comparison of *Buchnera* between *orange* and *green* variants retrieved 206 genes differentially expressed (p-value<=0.05) with 101 more expressed in the *green* variant (49%). However, even on differentially expressed genes, we could not find any signal of a link between differential methylation and expression.

As methylation is known in bacteria to be part of the immune system [47, 48] by being used with restriction enzymes, we also checked whether methylation occurred more frequently on DNA sequences that could be targeted by 101 restriction enzymes (list in Supp Mat, Table S7) whose DNA target could include a CpG dinucleotide. However, no significant methylation enrichment was found on those sites.

This, combined to the global correlation between clonal variants, prompted us to look at average methylation patterns at a more global scale, and relate them to genome composition.

We investigated the distribution of average methylation along the *Buchnera* genome. We observed a clear hotspot of methylation (Fig. 5A). As this effect seems regional, we defined the hotspot as the region with an above than average methylation percentage on Fig. 5A, including 10 out of the 14 genes showing a significant methylation enrichment. The hotspot starts at the CDS 472 and is composed of 47 genes. GO analyses showed that these 47 genes encode proteins involved in the translation machinery with a strong presence of ribosomal proteins (Table 2). Little is known about the role of cytosine methylation on ribosomal DNA sequences in bacteria [47]. Cytosine methylation is however known to be involved in regulation of gene expression (reviewed in [48]); such a mechanism, if applicable in the methyltransferase lacking *Buchnera*, could then help to regulate the growth of the endosymbiontic *Buchnera* population. Another possible role for these sequence methylation would be to hide them from the aphid host immune system. Indeed it is known that, even if the aphid has a reduced immune system to accommodate its endosymbionts, these symbionts may provoke an immune response [49,50,51]. Moreover, naked CpG are motifs that can be detected by the immune system [52]: the methylation of ribosomal genes of relatively high CpG content in Buchnera could then be thought of as a way for protecting them and dampen the immune system activation. This hypothesis is yet highly speculative as, if the aphid genome contains most of the Toll pathway, some Toll receptors are missing [53].

**Fig 5:**
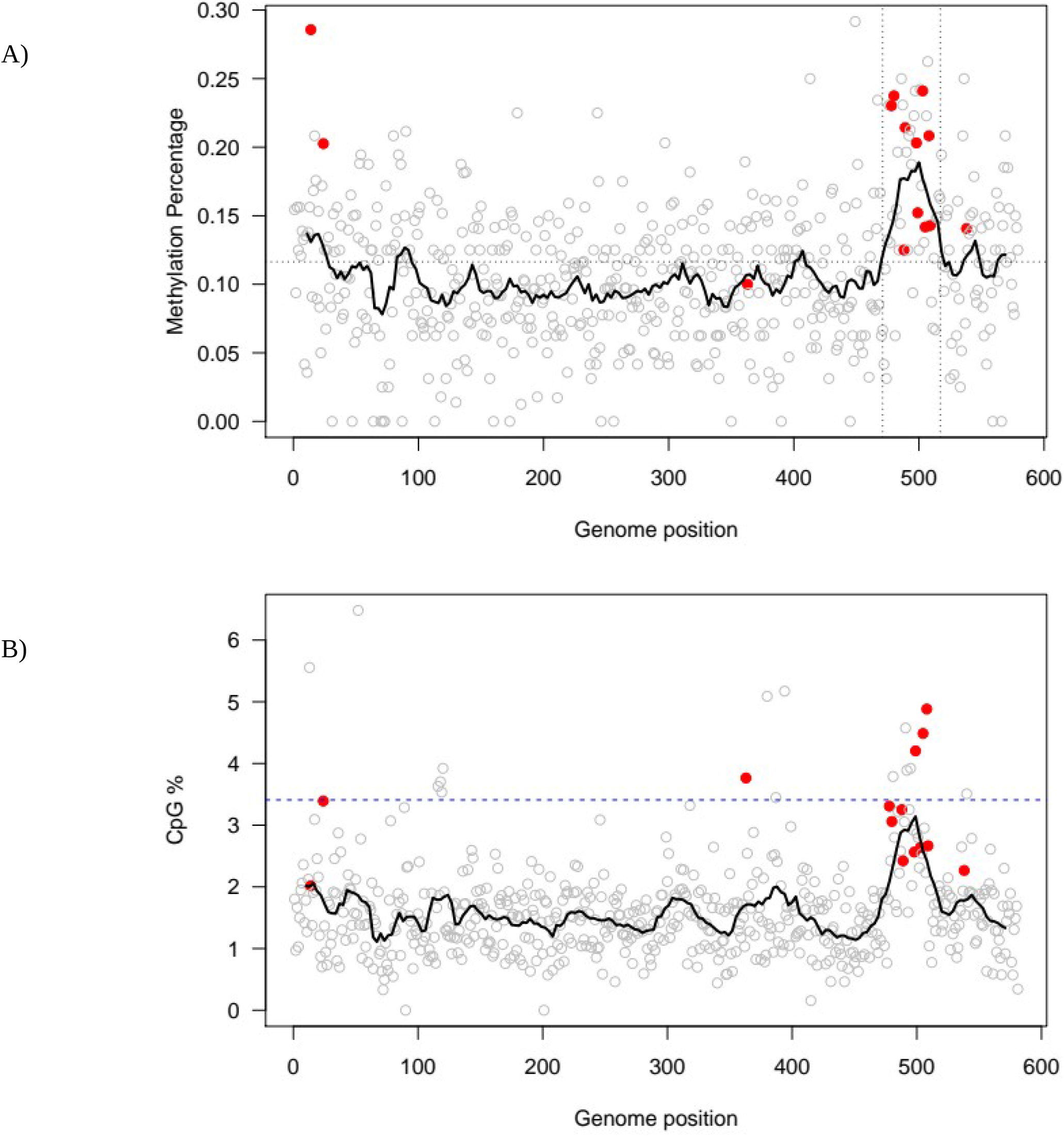
A) Methylation percentage in the Buchnera genome. x-axis represents the genome with 581 genes. y-axis is the methylation percentage (from 0 to 1). Each grey dot is a gene; red dots are genes that show a significant methylation enrichment – some genes with a high percentage of methylation are not enriched because they have few CpG sites. The black line is a sliding window average on 19 genes to emphasize regional characteristics. The horizontal dotted lines indicate the average methylation (∼11%), the vertical ones delimitate the hotspot. B) CpG dicnucleotide percentage in the Buchnera genome. x-axis represents the genome with 581 genes. y-axis is the CpG content. Each grey dot is a gene; red dots are genes that show a significant methylation enrichment. The black line is a sliding window average on 19 genes to emphasize regional characteristics. The blue dashed line represent the aphid host average CpG dinucleotide frequency.

**Table 2:**
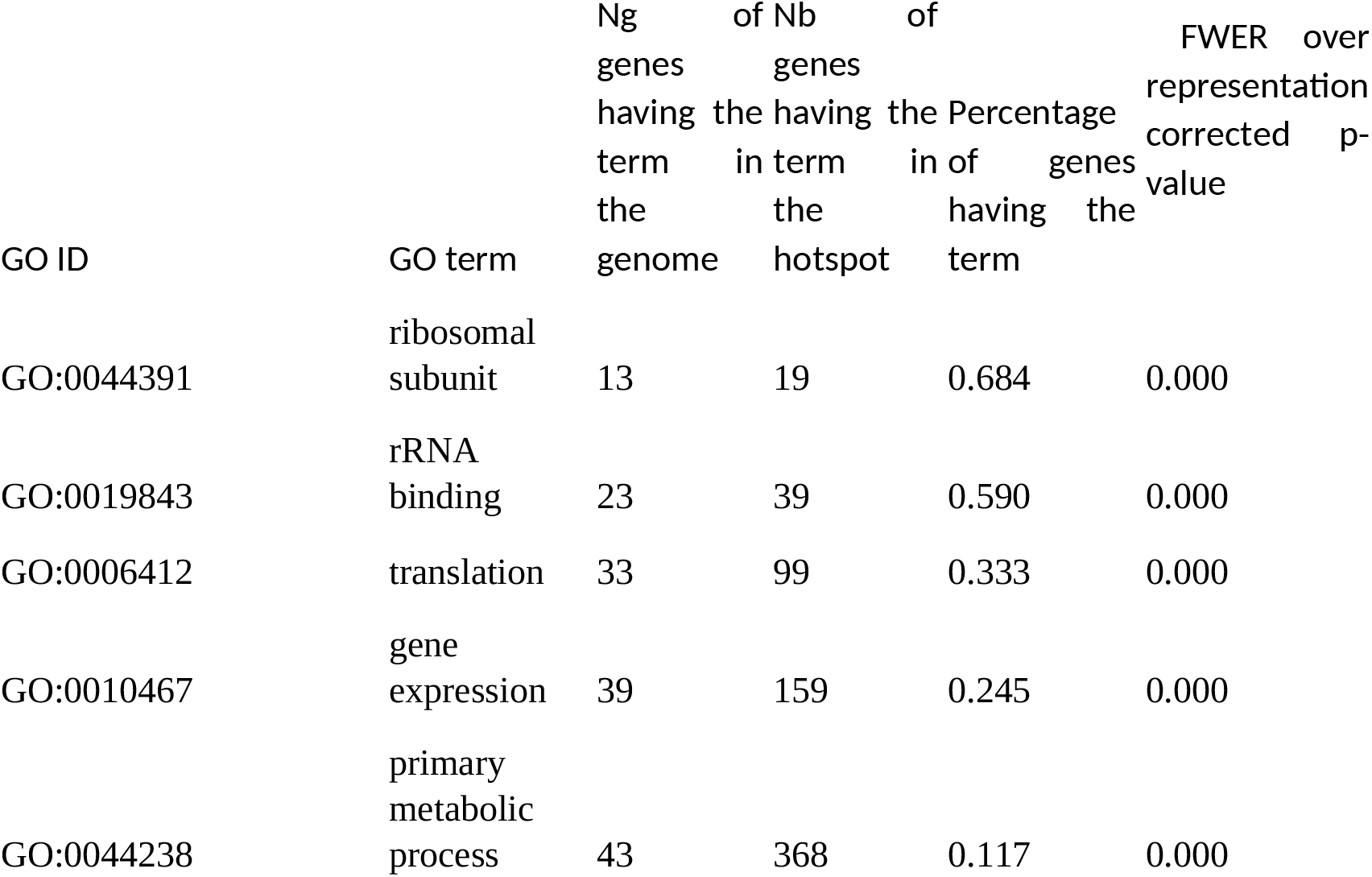
Main GO terms enriched in the Buchnera hotspot; a full table of all significantly enriched terms is given in Supp. Mat (Supp Table S8).

As a negative relation was found in the aphid host between methylation and CpG enrichment, we looked into CpG distribution along the *Buchnera* genome. We first observe a CpG dinucleotide frequency increase in the hotspot region (Fig. 5B), up to a value close to the one observed in the aphid genome. Moreover, a CpG enrichment is detected among hypermethylated genes (t-test between enrichment z-scores, p< 7. 10^*-5*^), meaning that the increase in CpG dinucleotide frequency in the hotspot is not a trivial consequence of regional base composition change, but that CpG dinucleotide are somehow selected for in this region. This is opposite to what was observed in the aphid host, and points a minima at the absence of counter-selection against CpG dinucleotide observed in the aphid host. This could be related to the absence of methyltransferase system in Buchnera, and then an hypothetical lack of role of the DNA methylation: methylation caused hypothetically by an aphid methyltrasferase leakage in Buchnera cells could not be efficiently used by eh endosymbiont, lacking the machinery to exploit and regulate such phenomenon. Going further, if indeed there is a positive selection on CpG dinucleotides in this region, as our data suggest, this could hint at a functional, yet unknown, role of the CpG dinucleotides in this region, maybe related to methylation.

## Discussion

The *A. pisum* genome wide methylation at CpG sites represents 3%, which appears low by comparison to mammals (80%) or honeybees (30%) [8,11,12,30,54]. However, this rate is higher than the one in the cotton bollworm (0.86%) [55], and these methylation are concentrated on gen sequences, where they span around 9% of the sequences. Although a low level of methylation mark is observed, aphids are equipped with a functional DNA methylation system and an extended set of chromatin modifying genes to process robust epigenetic regulation. Amazingly, *Buchnera* reduced genome shows an absence of genes coding DNA methylation and methyl sensitive restriction enzymes [56]. However we show that *Buchnera* from the two host variants presents methylated genes with a common hotspot. Methyl CpG are present at a frequency of around 10% in endosymbiont genome even though it is deprived of DNA methylase genes. However, the methylation in the endosymbiont genome appears robust in accordance with the levels found in E coli, close to 1% of all cytosines methylated by *dcm* [57]. Although there are solid correlation between a subset of aphid genes that appears heavily methylated and expressed over the average level of the full transcriptome, the differentially expressed genes appeared not related to a methylation state. More likely and as reported in many studies in other models, the methylation in aphid might shape the 3D architecture of chromosomes acting secondarily on the silencing or activation of genomic zones [8]. Interestingly, hot spots of methylation were consistently observed indicating a probable regulatory mechanism in regional chromosomal context.

On the side of developmental stages, the granddaughters of a female aphid (viviparous mother) are already developing within the daughters inside the ovarioles of the mother. Bacteriocytes that are large polynucleated cells filled up with *Buchnera* endosymbiont are transferred laterally from the mother to the ovarioles. These telescoping generations in clonality context constitute a powerful system to transfer trans-generationally multiple environmental signals [28,29]. In *Buchnera* most of genes for DNA repair and recombination like RecA, an essential component for the homologous recombination, are absent. Most of genes for the biosynthesis of the LPS components (lipo-poly-sacharides) that makes the cellular wall of bacteria are missing [37]. Surprisingly, most of genes responsible for phospholipid biosynthesis, indispensable for the formation of the membrane lipid bilayer, are also absent, except that for cardiolipin synthetase [37]. *Buchnera* seems to have delegated its membrane bilayer formation to the host. Our data suggest that *Buchnera* either imports methylases without identified signal motifs or synthesizes them in situ from host messenger, re enforcing the view that *Buchnera* is still far to evolve towards an organelle as mitochondria. The cell surface of *Buchnera* is known to be covered with hundreds of hook basal body made of flagellar proteins that have drifted putatively towards an import/export function helping to integrate the endosymbiont as an organelle [58]. Very little is still known about these structures in transport function. The canonical three-membrane system have been described for *Buchnera* endosymbionts sheltered in aphid bacteriocytes [56, 59]. Interestingly the symbiosomal membrane appears to form vesicles by membrane invagination, allowing to trap cytosol components for the *Buchnera* benefits. The dynamic of these structures are still unkown. Electron microscopy images reveals the importance of vesicle synthesis and trafficking inside the bacteriocytes. These vesicles tether the bacteria membrane and the host endoplasmic reticulum [56]. These cellular phenomenon have been corroborated by the high level of transcripts related of a Rab GTPase and vacuolar ATPase genes considered as markers of vesicle functioning [56,59,60]. Although the genesis of these vesicles is little studied and understood, we might hypothesize that bacteriocyte cytosolic molecules are trapped in a vesicle by membrane invagination and transferred in bacteria. Reticulum endoplasmic and bacteria could be also in a physical continuum, exchanging molecules. Theoretically, vesicles with bacteriocyte nuclear content could tether the endosymbiont membrane and shed aphid RNA messengers into *Buchnera* cytosol. These vesicles may mediate directional trafficking of molecules through cell surface in parallel with the one operating through the reticulum endoplasmic and Golgi. Dense vesicle presence within the bacteriocytes argues in favor of enhanced trafficking of bigger size molecules than metabolites, which might hint that RNA messengers and protein are exchanged between the insect body an endosymbiont. The bacteriocyte methylase gene products could be transferred into *Buchnera* by a passive diffusion mechanism which should be different of the known translocation system common to other organelles like mitochondria and chloroplasts. The unique case of one gene located in the aphid host genome and the gene product exclusively targeted in endosymbiont (RIpA4 gene) obviously does not apply to methylases that are deprived of signal peptide [61]. We might anticipate that the fragile and porous cellular membrane of *Buchnera* likely facilitates the transfer of macromolecules in both direction, accomplishing cross talk regulation to adapt to environmental changes. Presently no evidence support that aphid methylase could be addressed by signal peptide and targeting motif into the endosymbiont. Unknown possibilities can exist to transfer molecules in both directions between aphid host and endosymbiont in bacteriocyte context. Because vesicles have been shown on electron microscopy photographies, we might suggest that micro-vesicles (MV) are possible vehicles for proteins and mRNA. Micro vesicles have been abundantly described in host invading pathogens like *Leishmania, Cryptococcus*, and *Trypanosoma* and have been described to incorporate peptides, proteins, mRNAs and retrotransposons and lipids [60]. Evidence supports the concept of material delivery on long distance from cell to cell to induce metabolic stimulation or reprogramming of cells upon infection [60]. Diffusion of large size molecules between bacteriocyte and *Buchnera* cytosols for proteins that do not have a recognition motifs for targeting is suggested by few observations: a LD-carboxypeptidase and N-acetylmuramoyl-L-alanine amidase like genes, both required for bacterial peptidoglycan synthesis and/or recycling, are present in the aphid genome and absent in *Buchnera* genome, which supposes protein or mRNA transfer into the bacterial endosymbiont. Glutamate racemase which is responsible for converting L-glutamate to D-glutamate, an essential component of peptidoglycan biosynthesis, is absent in *Buchnera* genomes originated from few species of aphid such as C*inara confinis*, C*inara pseudotaxifoliae, Cinara tujafilina, Cinara cedri, Diuraphis noxia, Tuberolachnus solignus, Aphis glycines*, and *Schlechtendalia chinensis*, but present in *Buchnera* of many other aphid species. Both LD-carboxy peptidase and Glutamate racemase are cytosolic proteins without signal peptide. Furthermore, the nuclear proteins: toposimerase (parC) and gyrase A; essential enzymes for chromosome segregation and maintenance of chromosomes in an under-wound state respectively are absent in *Buchnera* of the aphids species genomes: *Cinara confinis, Cinara pseudotaxifoliae* and *Cinara BC ifornacula*. This suggests an import of homologs from these species host. Finally, Glutamoyl tRNA synthase (gltx), Aspartyl tRNA synthase (asps) and Proline tRNA synthase (prosS) are absent equally in C*inara confinis, Cinara pseudotaxifoliae* and *Cinara BC ifornacula*, which advocates for the transfer into *Buchnera* of either amino acid tRNA or the enzymes or their mRNAs. All these molecules are cytosolic without known motifs for targeting. All these elements argue in favor of a more complex dialogue and exchange of large size molecules between endosymbiont and host by the way of the symbiosomal vesicle fusion, emerging from invaginated symbiosomal membrane.

On the other hand, DNA methylation in organelles has been described despite the fact that they do not harbor methylases genes, which seems the case for the organelle evolving *Buchnera*. For instance, 5-methylcytosine were detected in chloroplasts and chromoplasts and some experiences demonstrate that DNA methylation suppresses transcription of photosynthesis genes in plastid [62]. On the other hand, chloroplast DNA of *Chlamydomonas reinhardtii* of maternal origin is heavily methylated [63]. Chloroplast genes from the maternal parent are transmitted to all progeny, whereas the corresponding alleles from the paternal parent are permanently lost. A strong correlation exists between maternal inheritance of chloroplast DNA and its hypermethylation mostly at CpG [63]. The arginine-rich region in the responsible methylase CrMET1 might act as signal peptide allowing its translocated in the organelles [63]. DNA methylation in the nucleus, mitochondrion and chloroplast in cultured *Sequoia sempervirens* (coast redwood) was described to orchestrate cell-fate plasticity [64]. In these few examples, organelles DNA don’t have methylase genes but use imported methylases for regulation purpose. On the other hand, ectopically expressed DNMT1-isoform3 was found localized into human mitochondria and methylate CpG sites in situ. In addition, overexpression of DNMT1-isoform3 was shown to orchestrate mitochondrial biological functions [65]. Flaws of methylation pattern at CpG sites in human mitochondrial genome was described associated with several physiological and pathological disorders [66,67]. Mitochondrial DNA methylation patterns and mitochondrial Dnmt3 levels are abnormal in skeletal muscle and spinal cord in presymptomatic ALS mice (amyotrophic lateral sclerosis (ALS)) [66,67].

Although by proteomic approach has shown no evidence of protein transfer from host to endosymbiont [68], our data obtained on two aphid variants in clonality context hints that some epigenetic dialogue in mutualist association between symbiont and host could take place.

## Materials and methods

### Maintenance of aphids

Aphids *Acyrthosiphon pisum* were maintained on *Vicia faba* plants, in an incubation room at 18°C, with a light/dark photoperiodicity of 16/8 hours. To obtain a green phenotype, aphids were raised at 9°C (see [69] for more details).

### WGBS analysis

The full sequencing of the genomes of aphid host and endosymbiont was performed after bisulfite treatment by the professor Lyko at Heildelberg (Germany). The DNA extraction was performed with the isolated ovarioles. This technique provides at the genome scale the precise location of methyl cytosine. All four samples (2 for orange and 2 for the green phenotype) were barcoded on a single sequencing lane, resulting in a strand-specific total sequencing coverage of 13x. The bisulfite conversion worked with >99.9% efficiency. For more information about the procedure we refer to the Illumina protocol [70] Sequencing was performed on a Hiseq 2000 [71].

### Bioinformatic library bisulfite treatment analysis

For each library, the Bismark aligner (version: v0.14.3) was then used to align the reads on a virtual concatenated genome generated from the Acyrthosiphon pisum (http://bipaa.genouest.org/is/aphidbase/acyrthosiphon_pisum/downloads/) and two symbionts, Buchnera aphidicola str. LSR1 (gb: ACFK01000001.1) and Candidatus Regiella insecticola (gb:CM000957.1) [72]. The methylation rate of each cytosine was analyzed using a house R script using the MethylKIT package. Alignments were conducted as described elsewhere [73].

### RNAseq Analysis

Total RNA was extracted according to the recommendation of the sequencing company Genewiz. Quality of RNA was checked by Agilent bioAnalyzer using the Agilent RNA 6000 Nanokit (Agilent). The sequencing was carried out by Genewiz according to the Illumina RNAseq sample preparation workflow. The successive steps are rRNA depletion, RNA fragmentation, random primer RT, dUTP incorporation, adenosine tailing, Y shaped adaptor ligation, dUTP strand degradation and finally PCR with barcode linked to adapters. The sequencing was performed based on a read length of 2×50pb. A First bioinformatic analysis was performed by Genewiz. From the generated Fastq files, sequences reads were trimmed to remove possible adapter sequences and low quality bases at ends with an error rate <0.05. Sequence reads shorter than 30 nucleotides were discarded. The remaining reads were mapped to the two reference genome using CLC genomics workbench (Buchnera Aphidicola str.APS (Acyrthosiphon pisum) genomic DNA, GCA_000009605.1_ASM960v1 and Acyrtosiphon pisum (Acyr_2.0, Assembly ID:GCA_000142985.2). Total gene hit counts and RPKM values were calculated for each gene (as well as the Q Score) and a comparison of gene expression between the two phenotypes was performed using DESeq2. Genes with adjusted p-value <0.05 and log 2 Fold change>1 were called as differentially expressed genes.

### GO enrichment analysis

The GO enrichment analysis is a statistical analysis of GO term frequency differences between two sets of sequences (for instance, one group of sequences coming from the RNA-seq experiment performed in this study and a reference available in public or published database). Blast2GO [74] was used for the pea aphid expression analysis. For the references, we chose the InterProscan results (v2.1) from AphidBase (a reference information system providing genomic resources for the study of aphids), created by BIPAA (Bioinformatics Platform for Agroecosystem Arthropods, from the Plant Health and Environment Division of INRA).

GO enrichment analyses were then performed using the GOFuncR package and corrected p-values reported are the FWER p-values.

### Validation of endosymbiont methylome by manual bisulfite PCR sequencing

Six genes from the top 10 methylated ensosymbiont genes were selected for validation. gDNA was extracted from green and orange aphid ovarioles, using the ISOLATE II Genomic DNA Kit (Bioline). We prepared 3 samples of 10 ovarioles for each genotype (green and orange). Ovarioles were grinded in lysis buffer incubated 3 hours with proteinase K. gDNA was treated with bisulfite using the Methyl Detector Bisulfite Modification Kit (Active Motif) and then purified. PCR were performed on 10 CDS found methylated in the two phenotypes (list in supplementary data). Primers were designed using Methprimer to specifically amplify bisulfite modified DNA (uracil instead of unmodified cytosine) (list in supplementary data). PCR fragments were purified using the QIA Quick PCR Purification Kit (Qiagen) and then ligated in the PCRTM2.1-TOPO using the TOPO TA Cloning Kit (Invitrogen). TOP10 competent cells were transformed with the fragment inserted vectors and spreaded on LB Agar Petri dishes on Kanamycine (50μg/ml) and Xgal (40mg/ml). The DNA fragments were collected individually by picking the white transformed colonies. For each genotype and each PCR, 5 colonies were sent for Sanger sequencing (GATC). Sequences were re analyzed based on C/U equivalence and unchanged methyl C. Only methylation on the two strands at CpG motifs were taken into account. The analysis of the sequences allowed us to verify on few selected genes the methylation rate obtained with WGBS (see Supp. Table S2 for detailed analysis of the manual validation of the WGBS data).

### Statistical analyses

All statistical analyses were done on R 3.6.3.

In aphid, the link between differential expression and differential methylation between green and orange strains was tested as follows: on genes begin differentially expressed between variants, a Spearman correlation test was done between the difference in methylation in green and orange variant, and the difference in expression.

In both aphid and Buchnera, sequences significantly hypermethylated were detected as those having a corrected p-value smaller than 0.05 when computing their occurrence probability with a Poisson distribution with mean the methylation average on genes in the organism considered. All 4 replicates (2 green and 2 orange) were pooled in this analysis. Multiple test p-value correction was done with Benjamini-Hochberg FDR.

CpG enrichment was computed using z-score in package seqinr [39], which accounts for sequence length and is not biased towards high scores for short sequences as is the rho statistic [75]. For both aphid and Buchnera, those z-scores were compared with a t-test between genes detected as hypermethylated and all other genes.

## Acknowledgements

We are very grateful to Jean Jacques Rémy, Minoo Rassazouldegan and Claude Pasquier for fruitful discussions that inspired the initial work. This work was supported by the ANR “blanc”, acronym: “Gustaile”, and the ANR “blanc”, acronym “methylclonome” and by the French Government (National Research Agency, ANR) through the LABEX SIGNALIFE program (reference # ANR-11-LABX-0028-01).

## Competing interests

The authors declare no competing interests

## Author contributions

M.C., M.D. A.R and M.B.B conceived the design of the experimental protocol and preparation of biological samples. S.A. did the validation of the WGBS/RNAseq data and the design/execution of the graphs related to WGBS and RNAseq datasheets. G.R. did the WGBS and initial analysis. MBB did statistical analyses. M.C.,M.D., M.B.B. and A.R. did to the manuscript writing.

## List of Supplementary Materials

**Table S1: Detailed results of Bismark analysis on the aphid genome**

**Table S2: List of genes identified as methyalated in [36] and comparison to i) the honeybee methylation status and ii) methylation status in our study**

**Table S3: Identification of the differentially expressed genes in aphid hosts and their methylation status**

Green and orange colored columns that indicates the log 2 fold changes correspond to the overexpressed genes in the *green* and *orange* phenotypes respectively. The methylation state of these genes is indicated.

**Table S4: list of hypermethylated genes in aphid host**

**Table S5: List of GO terms enriched among hypermethylated genes in aphid**.

**Table S6: Manual bisulfite sequencing performed on 6 genes found methylated in *Buchnera* by WGBS**

**Table S7: List of enzymes names and DNA motifs that were tested for being targeted by methylation in *Buchnera***.

**Table S8: List of GO terms enriched among hypermethylated genes in *Buchnera***.

## Notes

### Competing Interest Statement

The authors have declared no competing interest.

