## Supplementary material for "Methylation at CpG sites in genomes of aphid *Acyrtosiphon pisum* and its endosymbiont *Buchnera*": Supp Table S6

|  | **List of CDS used for manual bisulfite sequencing validation** |  |  |  |  |  |  |  |
| --- | --- | --- | --- | --- | --- | --- | --- | --- |
|  |  |  | Orange |  |  | Green |  |  |
| CDS | Dbxref | Product | Methylated reads  in pull down | Reads from manual bisulfite  sequencing in the gene | Reads from manual bisulfite  sequencing in the  promoter | Methylated reads  in pull down | Reads from manual bisulfite  sequencing in the gene | Reads from bisulfite  Manual sequencing  in the promoter |
| 17 | Genbank:WP_009873980.1 | molecular chaperone GroEL | 3 | 11 | 1 | 4 | 12 | 9 |
| 150 | Genbank:WP_009874115.1 | NADH dehydrogenase  subunit G | 3 | 3 | 12 | 4 | 14 | 15 |
| 362 | Genbank:WP_009874330.1 | DEAD/DEAH box family ATP  -dependent RNA helicase | 7 | 5 | 6 | 8 | 26 | 3 |
| 363 | Genbank:WP_009874331.1 | polyribonucleotide  nucleotidyltransferase | 3 | 4 | 2 | 4 | 11 | 6 |
| 507 | Genbank:WP_009874479.1 | 30S ribosomal protein S7 | 4 | 6 | 2 | 2 | 22 | 0 |
| 503 | Genbank:WP_009874475.1 | 50S ribosomal protein L3 | 3 | 9 | 3 | 1 | 4 | 0 |

| **List of primers used for the validation** | |  |
| --- | --- | --- |
| CDS | Forward primer | Reverse primer |
| 17 | 5' AGGAGAAAAATATTTTATTTAAAGT 3' | 5' AATAACATATAATACATCTTCAACAC 3' |
|  | 5' GTGTTGAAGATGTATTATATGTTATT 3' | 5' ACAAAAAATTCTTCACCAAAATTAAAA 3' |
| 150 | 5' TGGTTATTGTTATTTGTAAGATATGAT 3' | 5' AATAAATACTCCAATAAAACATAATTCTAT 3' |
| 362 | 5' ATTTTAATTTTATTATTATGATTT 3' | 5' ACTTTATCATCCTCTTTATT 3' |
|  | 5' TTGTAGAGTATAATTTATTTAATAAAGTAT 3' | 5' ATAAACTCAAAATCTTATATTCAT 3' |
| 363 | 5' GAATTAATGTGTTTTAGGTAAAATATTAGT 3' | 5' AAACAAAATAAAATTAAAAAATTAAATAAT 3' |
| 507 | 5' ATTGTTGTTTTTTTATTTTTTAAAGTATTA 3' | 5' CAACATATCAAATTCCTATTAAAATTC 3' |
| 503 | 5' TGTTTAAATTTTAATAGGTTGGTTTAA 3' | 5' ATCCCATAAAATACCTAATTCTAT 3' |

| **Results of manual bisulfite sequencing** |  |  |  |  |  |
| --- | --- | --- | --- | --- | --- |
| CDS | Number of analyzed sequences (colonies) | Number of methylated fragments in *orange* samples | Number of methylated fragments in *green* samples | Number of methylated sites | Percentage of methylated fragments |
| 17 | 30 | 2 | 0 | 7 | 6,67% |
|  | 30 | 1 | 1 | 2 | 6,67% |
| 150 | 30 | 0 | 0 | 0 | 0% |
| 362 | 28 | 2 | 1 | 3 | 10,71% |
|  | 28 | 3 | 0 | 3 | 10,71% |
| 363 | 30 | 1 | 1 | 3 | 6,67% |
| 507 | 30 | 1 | 3 | 5 | 13,33% |
| 503 | 30 | 1 | 1 | 2 | 13,33% |
