## Supplementary material for "Methylation at CpG sites in genomes of aphid *Acyrtosiphon pisum* and its endosymbiont *Buchnera*": Supp Table S3

| **Gene ID** | **Methylation percentage (orange)** | **Methylation percentage (green)** | **Mean FPKM in orange sample** | **SD of FPKM in orange sample** | **Mean FPKM in green sample** | **SD of FPKM in green sample** | **log2 Fold Change** | **Corrected p-value** | **Gene product** |
| --- | --- | --- | --- | --- | --- | --- | --- | --- | --- |
| ACYPI49688-RA | 0.00 | 0.01 | 1.42 | 0.36 | 259.37 | 90.85 | 5.98 | 8.2E-153 | myb-like protein P |
| ACYPI49694-RA | 0.00 | 0.03 | 0.81 | 0.34 | 255.85 | 108.46 | 5.88 | 4.7E-126 | putative uncharacterized protein DDB_G0282499 |
| ACYPI24862-RA | 0.67 | 0.78 | 0.33 | 0.09 | 19.93 | 5.42 | 4.49 | 6.8E-87 | alpha-tubulin N-acetyltransferase 1-like |
| ACYPI49687-RA | 0.00 | 0.01 | 0.55 | 0.22 | 36.44 | 19.49 | 3.90 | 3.7E-49 | uncharacterized LOC100574116 |
| ACYPI35620-RA | 0.00 | 0.00 | 1.10 | 0.49 | 27.27 | 4.27 | 3.75 | 2.9E-61 | uncharacterized LOC107883691 |
| ACYPI55671-RA | 0.00 | 0.00 | 1.71 | 0.07 | 36.32 | 4.01 | 3.73 | 1.1E-88 | uncharacterized LOC100575061 |
| ACYPI003728-RA | 1.00 | 0.75 | 0.44 | 0.01 | 12.29 | 4.22 | 3.65 | 1.6E-57 | alpha-tubulin N-acetyltransferase 1 |
| ACYPI25078-RA | 0.11 | 0.26 | 0.61 | 0.10 | 16.24 | 5.98 | 3.61 | 5.8E-55 | alpha-tubulin N-acetyltransferase 1-like |
| ACYPI35623-RA | 0.00 | 0.00 | 6.67 | 1.21 | 76.02 | 5.22 | 3.13 | 2.4E-57 | uncharacterized LOC100570351 |
| ACYPI009811-RA | 0.00 | 0.02 | 2.79 | 0.85 | 25.12 | 6.94 | 2.39 | 9.2E-34 | L-xylulose reductase |
| ACYPI071246-RA | 0.14 | 0.18 | 15.45 | 1.38 | 83.63 | 8.75 | 2.39 | 4.9E-56 | uncharacterized LOC100570586 |
| ACYPI49696-RA | 0.01 | 0.01 | 0.06 | 0.03 | 63.82 | 35.63 | 2.22 | 1.7E-15 | uncharacterized LOC100573859 |
| ACYPI009380-RA | 0.05 | 0.07 | 0.30 | 0.17 | 4.10 | 2.38 | 2.20 | 2.0E-15 | heat shock protein 83-like |
| ACYPI007260-RA | 0.00 | 0.01 | 2.00 | 0.35 | 9.43 | 0.95 | 2.15 | 4.7E-46 | plasma membrane calcium-transporting ATPase 1-like |
| ACYPI40829-RA | 0.00 | 0.01 | 0.77 | 0.13 | 6.66 | 4.13 | 2.04 | 4.6E-15 | uncharacterized LOC107884265 |
| ACYPI005472-RA | 0.00 | 0.01 | 5.21 | 1.38 | 24.61 | 3.82 | 1.94 | 5.0E-21 | high affinity cationic amino acid transporter 1 |
| ACYPI006423-RA | 0.01 | 0.01 | 12.08 | 0.67 | 13.65 | 2.30 | 1.90 | 2.7E-27 | protein NDRG1 |
| ACYPI004439-RA | 0.20 | 0.38 | 0.15 | 0.08 | 1.89 | 1.29 | 1.61 | 1.1E-08 | staphylococcal nuclease domain-containing protein 1-like |
| ACYPI082225-RA | 0.11 | 0.05 | 0.06 | 0.02 | 1.13 | 0.22 | 1.60 | 1.2E-08 | uncharacterized LOC100571498 |
| ACYPI35925-RA | 0.00 | 0.00 | 0.27 | 0.35 | 6.42 | 5.07 | 1.59 | 1.4E-08 | uncharacterized LOC100573363 |
| ACYPI51835-RA | 0.00 | 0.00 | 0.49 | 0.31 | 4.13 | 3.30 | 1.56 | 2.8E-08 | U3 small nucleolar ribonucleoprotein protein MPP10-like |
| ACYPI071401-RA | 0.02 | 0.01 | 2.24 | 1.13 | 8.50 | 1.16 | 1.52 | 2.3E-09 | uncharacterized LOC103307921 |
| ACYPI52083-RA | 0.00 | 0.01 | 0.18 | 0.21 | 4.37 | 2.83 | 1.52 | 7.2E-08 | uncharacterized LOC100570389 |
| ACYPI006920-RA | 0.00 | 0.00 | 4.05 | 1.04 | 16.26 | 6.36 | 1.49 | 2.4E-09 | uncharacterized LOC100166012 |
| ACYPI005423-RA | 0.00 | 0.05 | 0.22 | 0.12 | 1.58 | 0.75 | 1.46 | 2.1E-07 | protein aubergine-like |
| ACYPI009069-RA | 0.12 | 0.08 | 0.27 | 0.11 | 1.60 | 0.90 | 1.31 | 5.8E-07 | baculoviral IAP repeat-containing protein 7-B-like |
| ACYPI21300-RA | 0.01 | 0.01 | 1.04 | 0.41 | 4.04 | 1.92 | 1.28 | 2.4E-06 | uncharacterized LOC107884750 |
| ACYPI24137-RA | 0.39 | 0.34 | 0.30 | 0.14 | 2.56 | 1.63 | 1.24 | 1.0E-05 | uncharacterized LOC100570739 |
| ACYPI009213-RA | 0.00 | 0.00 | 0.10 | 0.09 | 2.09 | 1.45 | 1.24 | 6.4E-06 | peroxidase-like |
| ACYPI000544-RA | 0.00 | 0.00 | 0.66 | 0.21 | 1.86 | 0.46 | 1.23 | 8.4E-08 | fatty acid synthase-like |
| ACYPI41425-RA | 0.00 | 0.00 | 0.20 | 0.16 | 1.74 | 0.95 | 1.18 | 2.6E-05 | uncharacterized LOC100571655 |
| ACYPI007677-RA | 0.17 | 0.24 | 0.31 | 0.40 | 3.82 | 2.68 | 1.18 | 1.5E-05 | calreticulin-like |
| ACYPI005710-RA | 0.00 | 0.00 | 0.24 | 0.08 | 1.37 | 0.98 | 1.17 | 3.1E-05 | polyadenylate-binding protein 4-like |
| ACYPI41175-RA | 1.00 | 0.32 | 0.00 | 0.00 | 0.02 | 0.03 | 1.16 | 2.7E-05 | uncharacterized LOC100574043 |
| ACYPI007433-RA | 0.00 | 0.01 | 5.87 | 1.78 | 15.22 | 5.49 | 1.15 | 2.1E-06 | uncharacterized LOC100166574 |
| ACYPI007976-RA | 0.00 | 0.01 | 1.34 | 1.57 | 32.58 | 14.93 | 1.15 | 1.4E-05 | uncharacterized LOC100167162 |
| ACYPI56240-RA | 0.00 | 0.00 | 0.91 | 0.24 | 3.90 | 2.20 | 1.14 | 3.2E-05 | uncharacterized LOC100575341 |
| ACYPI56613-RA | 1.00 | 1.00 | 7.37 | 2.09 | 17.86 | 1.22 | 1.14 | 2.8E-07 | gamma-glutamyl hydrolase A-like |
| ACYPI002298-RA | 0.00 | 0.00 | 16.37 | 5.46 | 37.76 | 10.44 | 1.14 | 3.5E-07 | trehalase |
| ACYPI26793-RA | 0.00 | 0.01 | 1.15 | 0.29 | 3.62 | 0.67 | 1.13 | 7.8E-06 | poly [ADP-ribose] polymerase 12-like |
| ACYPI005750-RA | 0.00 | 0.01 | 1.08 | 0.70 | 4.59 | 3.34 | 1.13 | 5.9E-05 | RNA-binding protein 14 |
| ACYPI47418-RA | 0.08 | 0.00 | 0.45 | 0.14 | 2.00 | 0.90 | 1.13 | 5.6E-05 | uncharacterized LOC107883632 |
| ACYPI069505-RA | 0.50 | 0.56 | 0.20 | 0.12 | 1.33 | 0.52 | 1.12 | 5.7E-05 | chromobox protein homolog 5-like |
| ACYPI003374-RA | 0.50 | 0.21 | 14.41 | 1.94 | 30.58 | 3.30 | 1.12 | 2.8E-11 | uncharacterized LOC100162209 |
| ACYPI003317-RA | 0.57 | 0.48 | 3.75 | 0.30 | 7.84 | 1.19 | 1.12 | 1.5E-09 | ring canal kelch homolog |
| ACYPI47959-RA | 0.00 | 0.05 | 0.63 | 0.54 | 5.24 | 3.17 | 1.11 | 5.4E-05 | small ubiquitin-related modifier 3-like |
| ACYPI27192-RA | 0.64 | 0.63 | 7.91 | 0.88 | 16.88 | 2.97 | 1.10 | 3.8E-15 | ABC transporter F family member 4-like |
| ACYPI088675-RA | 0.64 | 0.63 | 7.91 | 0.88 | 16.88 | 2.97 | 1.10 | 3.8E-15 | ABC transporter F family member 4-like |
| ACYPI25704-RA | 0.51 | 0.45 | 0.06 | 0.05 | 0.44 | 0.25 | 1.09 | 1.0E-04 | uncharacterized LOC100573806 |
| ACYPI008324-RA | 0.43 | 0.60 | 0.13 | 0.06 | 1.25 | 0.57 | 1.09 | 9.4E-05 | UBX domain-containing protein 1-like |
| ACYPI53198-RA | 0.00 | 0.01 | 0.31 | 0.11 | 1.69 | 0.70 | 1.09 | 1.2E-06 | proton-coupled amino acid transporter 1-like |
| ACYPI008201-RA | 0.00 | 0.01 | 0.24 | 0.22 | 13.33 | 9.62 | 1.08 | 2.2E-05 | peroxidase-like |
| ACYPI002980-RA | 1.00 | 0.96 | 1.10 | 0.15 | 2.43 | 0.19 | 1.08 | 2.2E-07 | Acetyl-coenzyme A transporter 1-like |
| ACYPI006418-RA | 0.95 | 0.74 | 24.58 | 3.89 | 52.39 | 9.99 | 1.07 | 5.3E-09 | T-complex protein 1 subunit beta |
| ACYPI006978-RA | 0.86 | 0.89 | 0.65 | 0.07 | 1.56 | 0.08 | 1.07 | 9.4E-09 | cytoskeleton-associated protein 5 |
| ACYPI25268-RA | 0.00 | 0.00 | 0.47 | 0.07 | 1.37 | 0.37 | 1.07 | 8.1E-05 | plexin-B |
| ACYPI44794-RA | 0.07 | 0.20 | 1.22 | 0.18 | 3.14 | 1.21 | 1.07 | 2.7E-05 | small ubiquitin-related modifier-like |
| ACYPI006726-RA | 0.71 | 0.28 | 0.34 | 0.06 | 1.24 | 0.67 | 1.07 | 9.3E-05 | staphylococcal nuclease domain-containing protein 1-like |
| ACYPI56322-RA | 0.00 | 0.01 | 0.67 | 0.57 | 4.57 | 3.37 | 1.06 | 1.5E-04 | heterochromatin protein 1-like |
| ACYPI009024-RA | 0.00 | 0.00 | 3.08 | 1.66 | 10.25 | 6.07 | 1.05 | 1.7E-04 | esterase FE4-like |
| ACYPI007176-RA | 0.24 | 0.33 | 0.16 | 0.14 | 1.22 | 0.49 | 1.05 | 1.8E-04 | probable cleavage and polyadenylation specificity factor subunit 2 |
| ACYPI065075-RA | 0.00 | 0.00 | 0.18 | 0.10 | 1.41 | 0.87 | 1.05 | 1.6E-04 | MD-2-related lipid-recognition protein-like |
| ACYPI067858-RA | 0.38 | 0.44 | 14.00 | 0.82 | 28.05 | 2.86 | 1.05 | 2.3E-11 | acyl-CoA Delta(11) desaturase |
| ACYPI009434-RA | 0.01 | 0.00 | 9.85 | 4.95 | 35.56 | 13.82 | 1.04 | 2.3E-04 | zinc transporter ZIP1-like |
| ACYPI002182-RA | 0.02 | 0.01 | 47.91 | 11.29 | 103.28 | 42.28 | 1.03 | 5.9E-06 | late histone H2B.L4-like |
| ACYPI065029-RA | 0.01 | 0.00 | 0.26 | 0.26 | 7.34 | 4.92 | 1.02 | 1.7E-04 | uncharacterized LOC100575228 |
| ACYPI005279-RA | 0.00 | 0.03 | 0.43 | 0.06 | 2.32 | 1.46 | 1.02 | 2.8E-04 | small nuclear ribonucleoprotein-associated protein B-like |
| ACYPI20144-RA | 0.00 | 0.00 | 0.52 | 0.37 | 1.44 | 0.17 | 1.02 | 2.6E-04 | kelch-like protein 12 |
| ACYPI52071-RA | 0.56 | 0.51 | 0.52 | 0.15 | 2.08 | 0.88 | 1.01 | 3.2E-04 | uncharacterized LOC100575861 |
| ACYPI064994-RA | 0.43 | 0.41 | 0.69 | 0.22 | 1.83 | 0.55 | 1.01 | 1.5E-04 | transmembrane and ubiquitin-like domain-containing protein 1 |
| ACYPI086030-RA | 0.00 | 0.00 | 158.49 | 42.19 | 322.63 | 28.00 | 1.00 | 4.2E-07 | glycine-rich cell wall structural protein 1.8-like |
| ACYPI008524-RA | 0.05 | 0.00 | 3.05 | 0.87 | 7.71 | 3.66 | 1.00 | 1.8E-04 | cuticle protein 21-like |
| ACYPI34143-RA | 0.00 | 0.00 | 10.57 | 7.01 | 2.63 | 0.72 | -1.00 | 3.4E-04 | uncharacterized LOC100568855 |
| ACYPI42069-RA | 0.00 | 0.01 | 0.29 | 0.15 | 0.06 | 0.04 | -1.05 | 1.6E-04 | nose resistant to fluoxetine protein 6-like |
| ACYPI50492-RA | 0.00 | 0.00 | 4.46 | 2.46 | 1.13 | 0.17 | -1.06 | 1.3E-04 | uncharacterized LOC100569341 |
| ACYPI005811-RA | 0.00 | 0.00 | 5.34 | 1.53 | 1.84 | 0.37 | -1.08 | 5.1E-06 | uncharacterized LOC100164824 |
| ACYPI006623-RA | 0.03 | 0.01 | 1.82 | 0.12 | 0.59 | 0.08 | -1.09 | 1.7E-06 | cytochrome P450 306a1 |
| ACYPI32518-RA | 0.80 | 0.76 | 1.01 | 0.31 | 0.31 | 0.10 | -1.12 | 2.1E-05 | uncharacterized LOC100571902 |
| ACYPI50196-RA | 0.00 | 0.00 | 21.53 | 11.62 | 5.74 | 1.67 | -1.13 | 2.3E-05 | uncharacterized LOC100571908 |
| ACYPI086816-RA | 0.00 | 0.01 | 0.80 | 0.32 | 0.20 | 0.05 | -1.15 | 2.1E-05 | uncharacterized LOC100569659 |
| ACYPI45037-RA | 0.00 | 0.00 | 2.25 | 0.41 | 0.71 | 0.14 | -1.21 | 9.2E-08 | uncharacterized LOC107882419 |
| ACYPI001467-RA | 0.88 | 0.82 | 10.06 | 2.01 | 2.94 | 0.57 | -1.24 | 3.3E-11 | enoyl-CoA hydratase domain-containing protein 3, mitochondrial |
| ACYPI008446-RA | 0.00 | 0.00 | 12.39 | 6.45 | 2.51 | 0.81 | -1.30 | 1.8E-06 | ovarian fibroin-like substance-1-like |
| ACYPI42068-RA | 0.04 | 0.00 | 0.94 | 0.04 | 0.12 | 0.08 | -1.31 | 3.1E-06 | uncharacterized LOC107884316 |
| ACYPI47647-RA | 0.00 | 0.00 | 17.28 | 1.57 | 5.29 | 0.78 | -1.38 | 2.4E-16 | alpha-tocopherol transfer protein |
| ACYPI008428-RA | 0.01 | 0.00 | 5.11 | 0.72 | 1.32 | 0.42 | -1.38 | 1.1E-09 | glucose dehydrogenase [FAD, quinone] |
| ACYPI48465-RA | 0.00 | 0.00 | 6.13 | 3.46 | 0.66 | 0.04 | -1.42 | 1.5E-07 | uncharacterized LOC100571382 |
| ACYPI009883-RA | 0.00 | 0.00 | 47.95 | 17.38 | 12.10 | 2.57 | -1.47 | 6.3E-12 | carotene dehydrogenase |
| ACYPI21995-RA | 0.17 | 0.22 | 3.23 | 0.82 | 0.72 | 0.11 | -1.50 | 1.7E-13 | domeless 1 |
| ACYPI000807-RA | 0.00 | 0.00 | 21.01 | 3.38 | 4.94 | 0.39 | -1.64 | 1.8E-18 | facilitated trehalose transporter Tret1 |
| ACYPI31358-RA | 0.23 | 0.31 | 12.97 | 0.48 | 3.01 | 0.16 | -1.66 | 1.1E-19 | putative ankyrin repeat protein L59-like |
