## Supplementary material for "Methylation at CpG sites in genomes of aphid *Acyrtosiphon pisum* and its endosymbiont *Buchnera*": Supp Table S1

**Result of *bismark***

|  | Apisum_orange1 | Apisum_orange2 | Apisum_green1 | Apisum_green2 |
| --- | --- | --- | --- | --- |
| Sequence pairs analysed in total | 31697965 | 33396475 | 29894416 | 40149458 |
| Total number of Cytosine analysed | 539069496 | 573070931 | 551951134 | 747092555 |
| Total methylated Cytosine in CpG context  M | 3556438 | 399796 | 3858306 | 5103763 |
| Total methylated Cytosine in CHG context | 51908 | 56041 | 98944 | 74005 |
| Total methylated Cytosine in CHH context | 164227 | 184161 | 402068 | 241481 |
| Total unmethylated Cytosine in CpG context  U | 105815551 | 112256090 | 110173957 | 148634685 |
| Total unmethylated Cytosine in CHG context | 84678743 | 89436505 | 86087063 | 116915267 |
| Total unmethylated Cytosine in CHH context | 344802629 | 367140168 | 351330796 | 476123354 |
| Methylated cytosine in CpG context  = M/(M+U) | 3.3% | 3.4% | 3.4% | 3.3% |
| Methylated cytosine in CHG context | <0.1% | <0.1% | <0.1% | <0.1% |
| Methylated cytosine in CHH context | <0.1% | <0.1% | <0.1% | <0.1% |
